# Parasite intensity is driven by temperature in a wild bird

**DOI:** 10.1101/323311

**Authors:** Adèle Mennerat, Anne Charmantier, Philippe Perret, Sylvie Hurtrez-Boussès, Marcel M. Lambrechts

**Affiliations:** Department of Biological Sciences, University of Bergen, Norway; EDYSAN, Université de Picardie Jules Verne, CNRS, Amiens, France; CEFE, Univ Montpellier, CNRS, IRD, INRA, Montpellier SupAgro, EPHE, Montpellier, France; MIVEGEC, Univ Montpellier, CNRS, IRD, Montpellier, France; Département de Biologie Écologie, Faculté des Sciences, Univ Montpellier, Montpellier, France

**Keywords:** host-parasite, weather, climate, *Diptera*, *Protocalliphora falcozi*, *Protocalliphora azurea*, ectoparasite, blowfly, passerine bird, *Cyanistes caeruleus*

## Abstract

Increasing awareness that parasitism is an essential component of nearly all aspects of ecosystem functioning, as well as a driver of biodiversity, has led to rising interest in the consequences of climate change in terms of parasitism and disease spread. Yet empirical knowledge on the extent and ways in which climatic factors affect parasite prevalence and intensities remains scarce. In an 18-year, multi-site, correlative study we investigated the contributions of weather variables and other factors to spatio-temporal variation in infestation by blowfly parasitic larvae (*Protocalliphora* spp.) in nests of Corsican blue tits (*Cyanistes caeruleus*). We found that ambient temperature during the nestling stage is strongly and positively related to parasite load (number of parasites per chick), both across broods when controlling for year, and across years. In addition, annual mean parasite load also increased with minimal spring temperature, and decreased with increasing average temperature in the previous summer. There was no indication of a dependence of parasite dynamics on host dynamics in this system, likely due in part to the wide host range of blowflies that do not solely rely on blue tit hosts. This suggests a major effect of temperature during the blowfly life cycle, with potential implications for blowfly – host interactions across their geographical range as climate keeps warming up. Finally, given that ambient temperature increases throughout the breeding season and that blowflies negatively affect survival and recruitment of blue tits, these results also mean that parasites, along with caterpillar availability, can drive selection for breeding date in this system.

## Introduction

As global climate is very likely to keep warming up, it is of growing importance to understand how populations and communities respond to variations in temperature and precipitation. While milder, wetter winters are expected for Northern Europe, in the Mediterranean most of the expected climatic change will likely translate into warmer, drier summers (IPCC 2013). Increasing awareness that parasitism is an essential component of nearly all aspects of ecosystem functioning, as well as a driver of biodiversity (Hudson et al. 2006), has led to a rising interest in the consequences of climate change in terms of parasitism and disease spread (Harvell et al. 2002, Brooks and Hoberg 2007). Attempts were therefore made to predict the direction of change in disease prevalence in response to climate warming (e.g. Møller et al. 2013). However, it seems that no single scenario is to be expected given the dynamic nature of host-parasite interactions, the huge variation in parasite life histories, and the complexity of their effects at multiple levels within ecosystems (Mas-Coma et al. 2009, Rohr et al. 2011, Altizer et al. 2013). Therefore, predictions of the effects of climate change on infectious diseases need to be supported by detailed empirical knowledge acquired regionally in well-studied host-parasite systems (Hernandez et al. 2013, Roiz et al. 2014).

Understanding the population dynamics of parasites in relation to that of their hosts has been a central focus in disease ecology. In comparison, the direct influence of abiotic factors on parasites remains little studied, despite the fact that many parasites have in their life cycles at least one outside-host stage, during which they are exposed to environmental variability. In particular, ectoparasites that spend a significant part of their life cycle as free-living (*i.e.* away from their host) are most likely to be affected by weather conditions and climate changes (Hernandez et al. 2013, Rose et al. 2014, Charlier et al. 2016, Ogden and Lindsay 2016). For example developmental time, activity levels, and survival of ectoparasites often display bell-shaped responses to climate, *i.e.* are highest at intermediate temperature and moisture values (reviewed in Ogden and Lindsay 2016). Local weather fluctuations may therefore result in either increased or decreased ectoparasite intensities depending on the current position of populations relative to these optima (Stromberg 1997, Elderd and Reilly 2014, Eads and Hoogland 2016). In addition, invertebrates (including ectoparasites) may be differently affected by climatic variability depending on their biology, and in particular on their ability to take refuge in micro-habitats that can buffer adverse climatic conditions (e.g. Roiz et al. 2014, Ogden and Lindsay 2016).

Within nesting cavities, birds are in contact with invertebrates, including nest-dwelling ectoparasites that use avian nest material as habitat (Loye and Zuk 1991, Christe et al. 1994, Clayton and Moore 1997). The fitness costs of infestation as well as the fitness benefits of host defense traits are well documented in a wide range of bird species (e.g. Møller et al. 1990, Eeva and Nurmi 1994, Cantarero et al. 2013). This contrasts with the currently limited understanding of the factors driving the large temporal and spatial variation often found in ectoparasite intensities (Hurtrez-Boussès et al. 1999, Heeb et al. 2000, Dudaniec et al. 2007, Moreno et al. 2009). It has been suggested that nest ectoparasites might be influenced by weather, nest size or composition, or interactions with other invertebrate species within the nest microhabitat (Bennett and Whitworth 1991, Heeb et al. 1996, Remeš and Krist 2005, Kleindorfer and Dudaniec 2009, Moreno et al. 2009), yet the contributions of these factors relative to host factors remain unclear.

Among the best-studied nest ectoparasites in free-ranging birds are the haematophagous larvae of *Protocalliphora* blowflies that feed on bird nestlings’ blood (Owen and Ash 1955, Møller et al. 1990, Bennett and Whitworth 1991, Eeva and Nurmi 1994, Hurtrez-Boussès et al. 1997, Dawson et al. 2005, Remeš and Krist 2005, Cantarero et al. 2013). The first eggs deposited by adult *Protocalliphora* blowflies hatch from the beginning of the bird nestling stage onwards, and develop into three larval stages before pupating. Adult blowflies are free-living and likely overwinter in litter or in old nest materials (Matyukhin and Krivosheina 2008). The reported host spectrum of *Protocalliphora* is large amongst hole-nesting birds (Owen and Ash 1955, Jamriska et al. 2010). In blue tits (*Cyanistes caeruleus*), *Protocalliphora* blowflies have well-established detrimental effects on nestling growth and survival, resting time, aerobic capacity at fledging, and eventually post-fledging survival and recruitment (Merino and Potti 1996, Hurtrez-Boussès et al. 1997, Charmantier et al. 2004, Simon et al. 2004, 2005, Thomas et al. 2007). In addition, *Protocalliphora* abundance is positively related to bacterial loads measured on nestlings, suggesting higher risks of bacterial infection in nests heavily infested by *Protocalliphora* (Mennerat et al. 2009). *Protocalliphora* do not parasitize adult birds, but indirectly affect parental effort as reflected in increased feeding rates or increased investment into nest-sanitation behaviour (Hurtrez-Boussès et al. 2000, Bańbura et al. 2004). On the island of Corsica, blue tit nestlings are exposed to the highest *Protocalliphora* loads reported so far in European study sites, while other types of nest ectoparasites are rarely found (Hurtrez-Boussès et al. 1997, 1999, Mennerat et al. 2008).

In an 18-year, multi-site, correlative study, we investigated how *Protocalliphora* infestation intensities varied both within and across years, with the main objective to understand the relative contributions of climatic and other factors to spatio-temporal variation in parasite intensities. We more specifically explored within-year variability in relation to host life history, nest characteristics and weather during the nestling stage, while accounting for spatial variation. We also explored how inter-annual variability in mean parasite intensities relates to host dynamics and life history, as well as to temperatures and rainfall during the summer, autumn, winter and spring preceding each breeding season.

## Methods

### Study sites and monitoring

Data from blue tit broods and uniquely ringed female breeders were obtained for years 1997-2014 from seven study plots located in two valleys on the island of Corsica: plots Avapessa, Arinelle, Feliceto, Filagna, Grassa and Muro in the Regino valley, and plot Pirio in the Fango valley) (Blondel 1985, Lambrechts et al. 2004). The broad-leaved deciduous oak *Quercus humilis* favouring the production of earlier and larger blue tit broods was the dominant tree species in plots Avapessa, Feliceto, and Muro. The evergreen oak *Q. ilex* was the dominant tree species in the other four study plots where blue tit broods are smaller and occur later in the season than in deciduous habitats (Lambrechts et al. 2004, Blondel et al. 2006). Breeding blue tits used either wood-concrete Schwegler B1 boxes (Schorndorf, Germany) or concrete boxes of similar dimensions (nest-chamber size of *ca.* 113 cm^2^).

Following basic protocols (Blondel et al. 2006) boxes were visited at least once a week to check the initiation and progress of nest construction, and determine the egg-laying date, number of eggs, and number of nestlings in the nest. Nest thickness, *i.e.* the vertical distance between the bottom floor and the top of the external nest wall (Hurtrez-Boussès et al. 1999), was measured either shortly before or during the egg-laying period. Given that all nestboxes in this study have similar internal diameters (12 cm), nest thickness is an appropriate proxy for nest volume, *i.e.* habitat size for nest ectoparasites.

Because nests were visited weekly at the end of the incubation stage, the onset of hatching was calculated based on the physical development of the nestlings at 0 to 6 days after hatching (Descamps et al. 2002). When nestlings were between nine and 15 days old, adult breeders were trapped inside the nestbox. The age of the female parent (yearling *vs* older) was determined either from the monitoring records for previously ringed birds, or by comparing the colour of the alula and primary wing coverts to that of greater wing coverts (Blondel et al. 2006). Parental female age may contribute to variation in parasite loads because nest sanitation behaviour (exclusively performed by females) may be affected by breeding experience (Hurtrez-Boussès et al. 2000, Banbura et al. 2001). Breeding attempts for which females could not be caught were not considered in this study.

### Meteorological data

Records of daily minimum temperature (°C), daily maximum temperature (°C), and daily rainfall (mm) were obtained from the meteorological station of Calvi in Corsica. This weather station is situated at a maximum of 20 km from each of the study plots and thus gives reliable information on regional meteorological variation (see also Grosbois et al. 2006). We averaged the daily minimum and maximum ambient temperature to estimate daily average ambient temperature. For each nest during the two weeks following hatching we calculated the average ambient temperature and the average amount of rainfall. This period corresponds to the time when *Protocalliphora* larvae develop in blue tit nests by intermittently feeding on nestling blood, while spending the rest of the time hidden amongst nest materials (Hurtrez-Boussès et al. 1999).

To explore the effect of inter-annual climatic variation on mean *Protocalliphora* intensities we used meteorological archives of minimal, maximal and average monthly temperatures, as well as monthly rainfall. We calculated average temperature and total rainfall over three-months periods corresponding to summer (June-August), autumn (September-November), winter (December-February), and spring (March-May) preceding each breeding season.

### Protocalliphora abundance

Previous studies (Hurtrez-Boussès et al. 1999) have shown that two blowfly species coexist in Corsican blue tit populations : *Protocalliphora azurea* (Fallén 1817) and *Protocalliphora falcozi* (Séguy 1928). Since it is impossible to morphologically distinguish between the two species at larval and pupal stages, we kept them pooled as *Protocalliphora.* Nests were collected 15 days post-hatching (*i.e.* one week before fledging), stored in hermetic plastic bags, and replaced in the nestbox by similar amounts of new nest material, mainly moss. In the laboratory, *Protocalliphora* larvae and pupae were carefully sorted out of the nest material and counted. Our counts included the total number of second-stage larvae, third-stage larvae and pupae (excluding first-stage larvae that are difficult to detect due to their small size), following the protocols presented in Hurtrez-Boussès et al. (1999), Heeb et al. (2000) and Mennerat et al. (2008; 2009). In some study years and sites (e.g. Mennerat et al. 2008; 2009), some nests were enclosed in cotton bags to facilitate their collection (without bag: 274 nests; with bag: 261 nests). Cotton bags were first inserted under blue tit nests around hatching time, so that the blue tit parents could habituate to the presence of the bag. A few days later the edge of the bag was pulled up to reach the same height as that of the nest.

### Statistical analyses

This study includes data from 535 broods covering an 18-year period (Table 1). Our study only focused on first-clutch broods for which the number of eggs and nestlings was not manipulated, nests that were not experimentally treated against parasites, and nests for which there was no evidence of predation (the main predator at these nestbox study sites is the green whip snake *Hierophus viridiflavus)*. We used the mean number of *Protocalliphora* larvae per nestling (*i.e.* the number of larvae and pupae found in a nest divided by the number of chicks present at time of sampling) as a measure of parasite load. All analyses were performed in the statistical programming environment *R 3.2.2* (http://r-project.org). Model validation was performed by visual inspection of residuals.

**Table 1.** Yearly fluctuations in the number (mean ± SD) of *Protocalliphora* larvae per blue tit nestling on Corsica from 1997 to 2014. Sample sizes (broods) are in parentheses.

| Year | Muro valley | Fango valley |
| --- | --- | --- |
| 1997 | - | 12.15 ± 7.01 (10) |
| 1998 | 6.98 ± 0.84 (2) | 10.38 ± 6.72 (8) |
| 1999 | 6.15 ± 1.21 (2) | 11.07 ± 4.53 (4) |
| 2000 | 3.66 ± 2.63 (6) | 13.79 ± 5.20 (5) |
| 2001 | 3.07 ± 3.48 (23) | 13.12 ± 5.51 (16) |
| 2002 | - | 9.54 ± 3.29 (3) |
| 2003 | - | 9.00 ± 3.59 (11) |
| 2004 | - | 5.78 ± 4.71 (19) |
| 2005 | 5.44 ± 3.65 (53) | 10.23 ± 5.41 (35) |
| 2006 | - | 12.44 ± 5.47 (6) |
| 2007 | 3.37 ± 2.97 (56) | - |
| 2008 | 5.68 ± 5.81 (52) | 15.31 ± 8.58 (23) |
| 2009 | 3.55 ± 2.73 (39) | 11.39 ± 6.36 (33) |
| 2010 | 2.94 ± 2.62 (29) | 8.11 ± 4.14 (35) |
| 2011 | - | 10.64 ± 6.71 (24) |
| 2012 | - | 12.90 ± 7.27 (13) |
| 2013 | - | 2.12 ± 1.49 (5) |
| 2014 | - | 12.08 ± 10.21 (23) |
| Total | 4.26 ± 3.93 (262) | 10.70 ± 6.77 (273) |

#### Within-year variation in parasite intensity

To investigate the relation between parasite intensity and current biotic and abiotic factors, we used linear mixed-effect models with log-transformed parasite load as a dependent variable (*lmer* from the lme4 package). We applied forward, AIC-based model selection starting with a set of models with nest thickness, cotton bag treatment (with *vs* without a cotton bag), egg-laying date (in Julian dates), female age, and weather during the nestling stage (average ambient temperature and rainfall) as explanatory variables. Year (n=18), valley (n=2), study site (n=7), nestbox identity (n=243), and female identity (n=385) were included as random effect factors.

#### Inter-annual variation in parasite intensity

We further investigated the inter-annual variation in mean parasite load in relation to weather during the summer, autumn, winter and spring preceding each breeding period. The study sites are located in two distinct valleys (Table 1) that differ markedly in a range of factors (Blondel et al. 2006). We used linear models with mean (yearly average calculated for each valley) *Protocalliphora* abundance per chick as a dependent variable. We applied forward, AIC-based model selection with an initial set of explanatory models with valley as a factor (to account for potential valley-specific relations between parasite intensities and weather) and yearly mean values of egg laying-date, ambient temperature during the nestling period, minimum, maximum, and average ambient temperature and total rainfall during each three-month period (*i.e.* season) preceding the breeding season when parasite intensities were sampled. Two measures of host performance in the previous year (average brood size and average fledgling mass) were also included in the initial set of variables, to account for a potential relation between parasite and host dynamics. Interactions between valley and all other covariates were also included. The three explanatory variables retained in the final model were not correlated.

## Results

### Within-year variation in parasite intensity

The final model for brood parasite intensity included only ambient temperature during the nestling stage and cotton bag treatment as explanatory variables, and its fit was further improved by adding a quadratic term for ambient temperature. Parasite load increased with mean ambient temperature (linear term: P < 10^−4^), with some degree of saturation at higher temperatures (quadratic term: P < 10^−4^; Table 2A; Figure 1).

**Figure 1.**
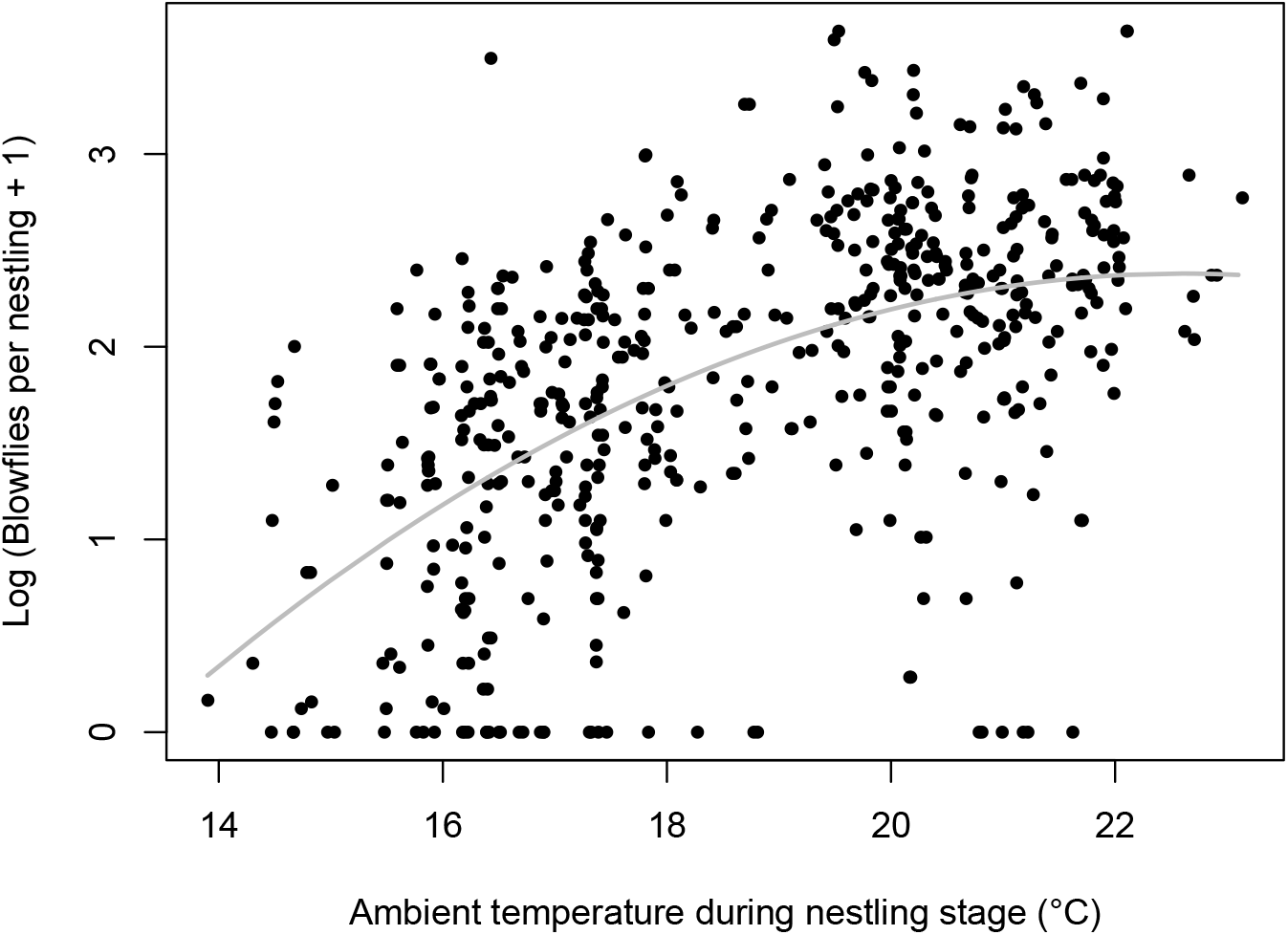
Across broods, parasite load increases with mean ambient temperature during the nestling stage (*i.e.* during the developmental period of *Protocalliphora* larvae) in nests of blue tits on Corsica. Each dot represents a brood. The grey curve shows predicted values from the final model (see Table 2).

**Table 2.**
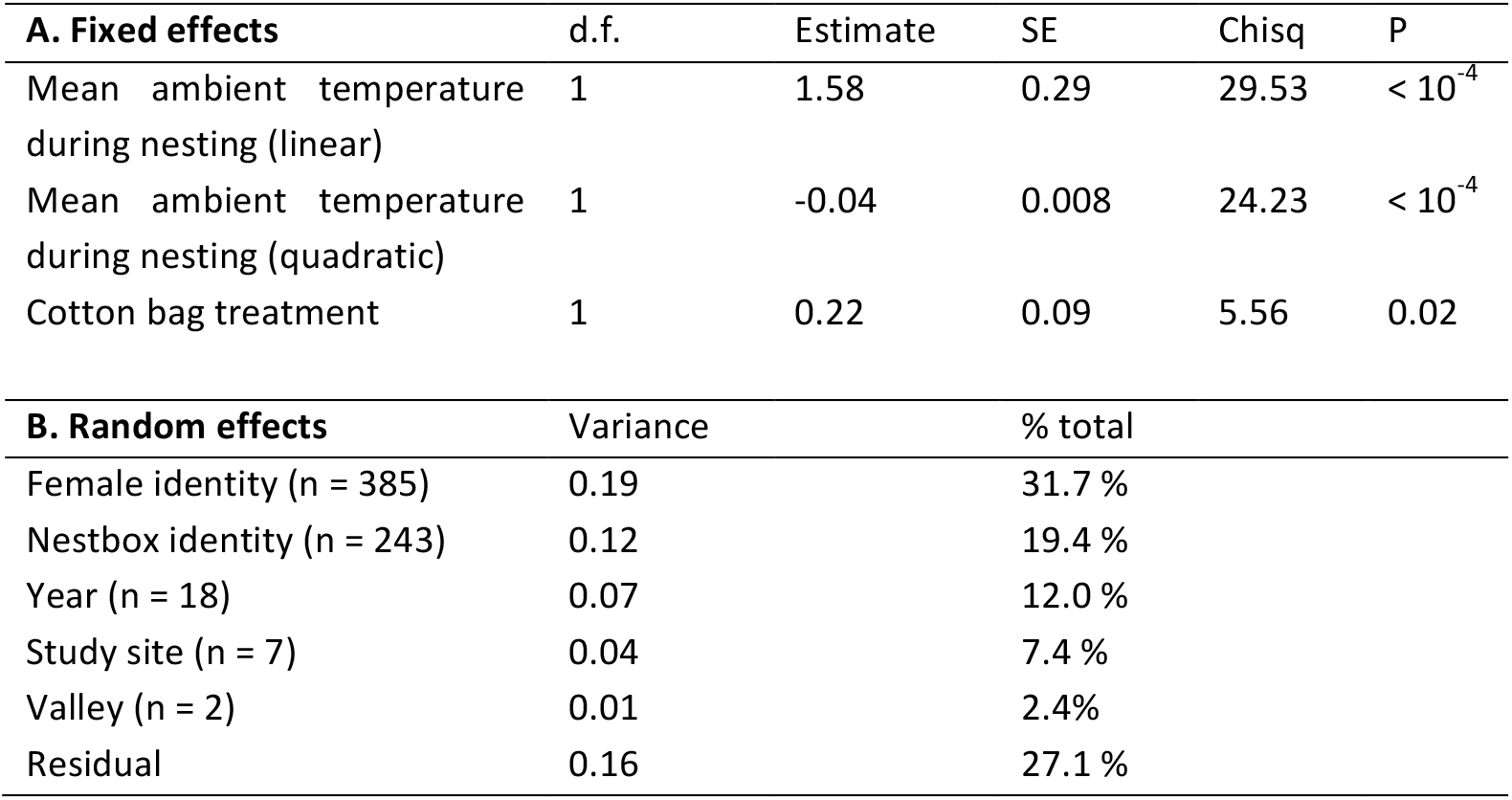
Within-year variation in parasite load per chick in blue tit nests infested by hematophageous *Protocalliphora* larvae, in relation to mean ambient temperature during the nestling stage (*i.e.* in the period when parasite larvae develop into blue tit nests) and cotton bag treatment (*i.e.* presence or absence of a cotton bag around the nest). Results from the final linear mixed effects model, obtained after forward selection from an initial set of models with nest thickness (*i.e.* height), cotton bag treatment, egg-laying date, female age, and average ambient temperature and rainfall during the nestling stage as explanatory variables. Year, valley, study site, nestbox identity, and female identity were included as random effect factors.

In addition, parasite load was higher in nests surrounded by a cotton bag than in nests collected without a cotton bag (P = 0.02). Among the random effects factors, more than half of the total variance was explained by female identity and nestbox identity together (female identity: 31.7%; nestbox identity: 19.4%; Table 2B). Year, study site and valley explained 12, 7.4% and 2.4% of the variance respectively.

### Inter-annual variation in parasite intensity

Across years, mean parasite load was strongly and positively correlated with ambient temperature during breeding (P < 10^−4^; Figure 2), and to a lesser extent negatively correlated with average temperature in the previous summer (P = 0.009; Figure 3) and positively correlated with minimal spring temperature (P = 0.03; Figure 4). No other variable was retained in the final model (Table 3). In particular, no significant difference was detected between the two valleys. Even after excluding the year with the warmest summer (see Figure S1C), model selection resulted in the same set of variables.

**Figure 2.**
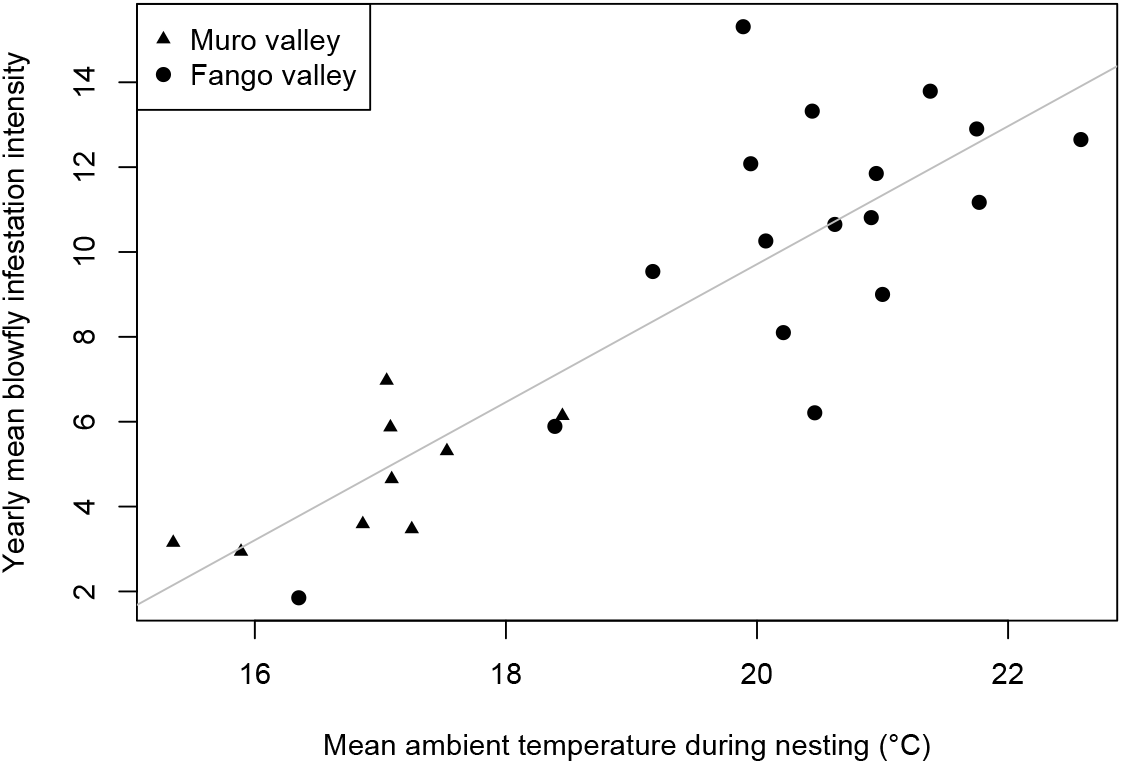
Across years, parasite load correlates positively with ambient temperature during the nestling stage (*i.e.* during the developmental period of *Protocalliphora* larvae) in nests of blue tits on Corsica. Dots represent yearly average values.

**Figure 3.**
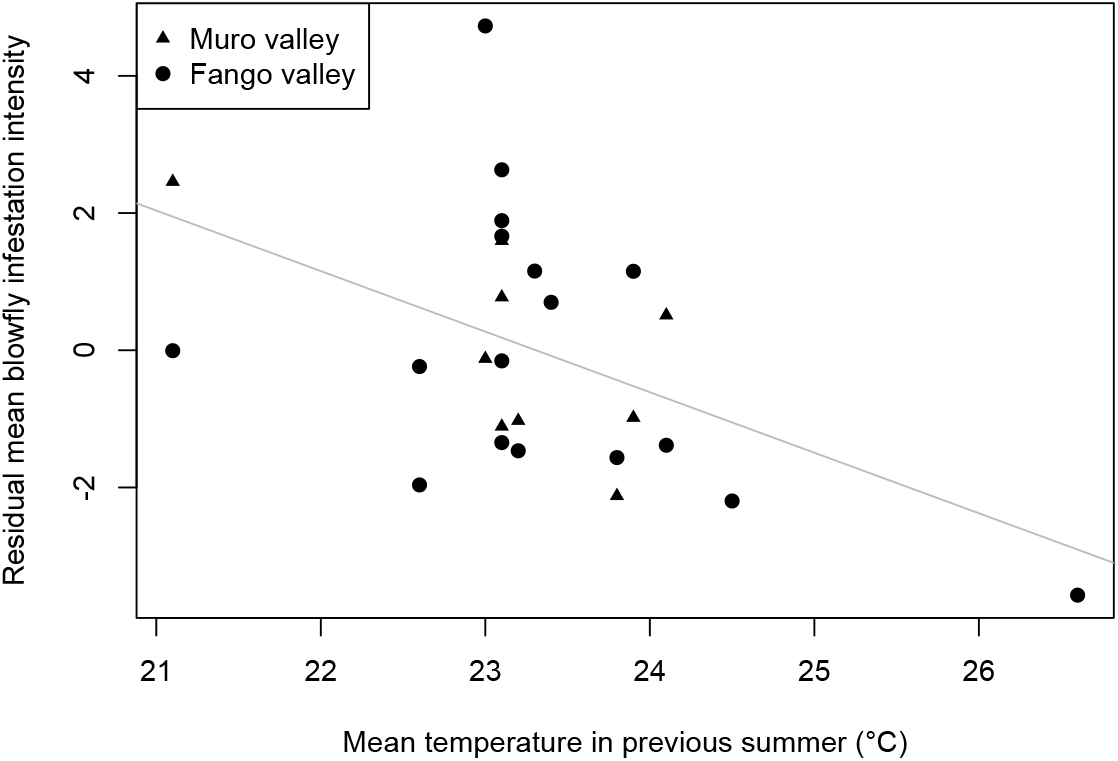
Across years, parasite load (corrected for ambient temperature during nesting) decreases with increasing mean temperatures in the previous summer (i.e. during the adult, free-living stage of *Protocalliphora* blowflies). Dots represent yearly average values.

**Figure 4.**
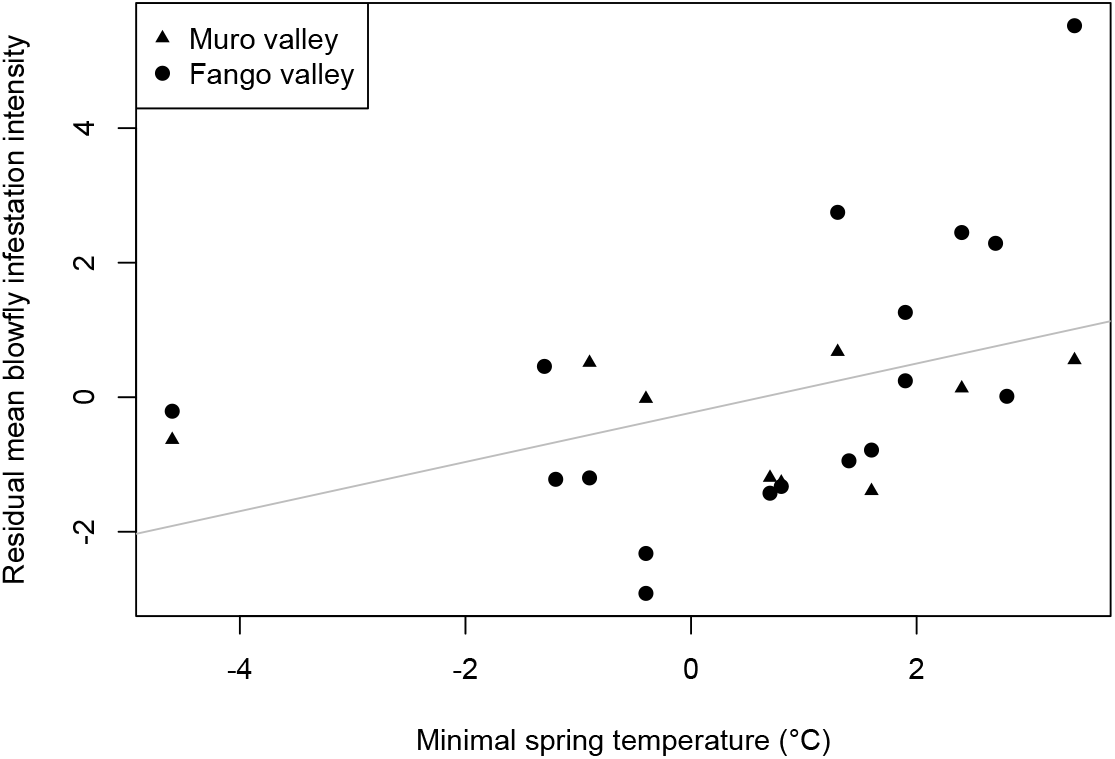
Across years, parasite load (corrected for ambient temperature during nesting) increases with increasing minimal temperatures in early spring (*i.e.* during the adult, free-living stage of *Protocalliphora* blow flies). Dots represent yearly average values.

**Table 3.** Inter-annual variation in mean parasite load per chick (yearly average) in nests of blue tits infested by hematophageous *Protocalliphora* larvae, in relation to yearly average values of mean temperature during nesting, mean temperature in the previous summer, and minimal spring temperature. Results from the final linear model, obtained after forward selection from an initial set of models with valley, egg laying-date, ambient temperature during the nestling period, minimum, maximum, and average ambient temperature and total rainfall during each preceding season as explanatory variables, as well as interactions between valley and all other covariates.

|  | d.f. | Estimate | SE | Sum of squares | P |
| --- | --- | --- | --- | --- | --- |
| Mean temperature during nesting | 1 | 1.60 | 0.17 | 272.1 | < 10<sup>-4</sup> |
| Mean temperature in previous summer | 1 | -0.89 | 0.34 | 20.40 | 0.009 |
| Minimal spring temperature | 1 | 0.37 | 0.17 | 14.32 | 0.03 |

## Discussion

This study reveals that ambient temperature during the nestling stage is strongly related to variation in parasite load, both across broods when controlling for year, and across years. In addition, annual mean parasite load (1) increased with minimal spring temperature, and (2) decreased with increasing average temperature in the previous summer. In this Mediterranean study system, parasite intensity thus appears to be primarily driven by ambient temperatures during spring and summer. Noticeably, as far as we could test, parasite dynamics did not relate to host dynamics. This might be explained by the fact that *Protocalliphora* blowflies have a broad host range and do not solely rely on blue tits, and that only the larval stage is parasitic. Finally, we found that both female identity and nestbox identity accounted for a relatively large amount of variance in brood parasite load. This finding that parasite loads are repeatable across females and nestboxes suggests that genetic and or maternal effects, as well as local environmental effects are important determinants of parasite abundances.

Our results suggest several ways in which ambient temperature may affect the life cycle of *Protocalliphora* blowflies. The main factor found to affect parasite load is ambient temperature during the larval stage (*i.e.* the nestling stage of blue tit hosts). In insects, temperature dependence of larval growth and survival has been studied extensively in the laboratory (overview in e.g. Chown and Nicolson 2004), yet surprisingly little in the field. Our study confirms, as observed in another host species the tree swallow *Tachycineta bicolor* (Dawson et al. 2005), that blowfly larval abundance is driven by temperature. In the tree swallow study, parasite abundance increased across the natural temperature range in a curvilinear way very similar to what we report here (see Figure 1), and decreased when temperatures were experimentally raised above natural levels (see also Castaño-Vázquez et al. 2018). As suggested in a recent study, a concomitant decrease in nest-dwelling ectoparasite loads could explain why experimental heat stress appears to have positive effects on blue tit fledglings (Andreasson et al. 2018). Our results here point to an optimal temperature for larval development around 23-25 °C and suggest that larval survival is significantly reduced below 20 °C.

We also found a negative effect of warmer summer temperatures on *Protocalliphora* loads in the following spring, possibly as a consequence of increased mortality of adult, free-living blowflies at high temperatures. While heat tolerance limits in adult *Protocalliphora* are currently unknown, in other dipteran species they appear to range from 30 °C and above (Feder et al. 1997, Berrigan et al. 2000, Chown and Nicolson 2004, Enriquez and Colinet 2017). Field studies of thermal stress in the willow beetle *Chrysomela aeneicolli*, which were carried out at a latitude close to that of our study sites, reveal that the temperatures measured on sunlit soil can approach 40 °C even when the maximum air temperature recorded by a nearby weather station did not exceed 26 °C. Considering that in our study area the maximal summer (air) temperature ranged from 34.0 up to 40.6 °C (average summer temperature 21.1 – 25.4 °C, see Figure S1), this means that adult *Protocalliphora* blowflies might in some years be exposed to temperatures above their thermal tolerance levels, resulting in high adult mortality. This, however, remains to be tested.

Minimal spring temperature (ranging from −4.6 °C to 3.4 °C, Figure S1) also seems to influence parasite loads, while minimal winter temperature does not (ranging from −4.2 °C to 1.9 °C). Given that all minimal spring temperatures were recorded in March, this might suggest that the end of the overwintering period for adult blowflies takes place during this month after (Matyukhin and Krivosheina 2008), and that blowflies are then vulnerable to cold-induced mortality. This explanation remains speculative, since little is currently known about the biology of blowflies in their free-living (*i.e.* adult) stage. More generally, the potential selective effect of temperature extremes on parasites in this system is an aspect that deserves further attention.

Because mean ambient temperature during the nestling stage increases linearly with egg-laying date, our results help explain why parasite loads are higher in late broods. They suggest temporal variation in the selection that parasites may impose on their hosts (Figure S2). In the focal study populations as in generally all temperate insectivorous birds, laying date is persistently under negative natural selection (Porlier et al. 2012) and the strength of selection is stronger during very warm springs (Marrot et al. 2017, 2018). This strong selection favouring early breeding females is always discussed in the context of the phenological mismatch between birds and their main caterpillar preys (e.g. van Noordwijk et al. 1995, Visser et al. 2006). The present study shows that nest ectoparasites may also be one of the drivers of selection on laying date. Despite high prevalences and intensities in Corsican populations of blue tits (Hurtrez-Boussès et al. 1997), no specific host behaviour was found to help preventing these parasites from accessing the nest (Mennerat et al. 2008) or removing them from nest material (Hurtrez-Boussès et al. 2000). It appears, however, that adult blue tits can compensate for some of the harmful effects of parasites by increasing rates of chick provisioning (Hurtrez-Boussès et al. 1998). The level of compensation depends on food availability (Simon et al. 2004), and, as a result, parasites and food interact to determine post-fledging survival and recruitment (path analysis, Thomas et al. 2007). Based on this, parasite-mediated selection is likely strongest in evergreen oak-dominated habitats, characterised by a combination of low caterpillar abundance and late breeding, and where parasite loads are highest. On the contrary in deciduous holm oak-dominated habitats, parasites might exert relatively weak selection on blue tits due to a combination of plentiful food, early breeding, and low parasite loads. The nature of traits in blue tits that could be under parasite-mediated selection has yet to be established, but breeding phenology appears as a likely candidate, as well as parental behaviours that improve nestling growth or survival under conditions of high parasite abundance (Hurtrez-Boussès et al. 1998, Mennerat et al. 2009). Our results also reveal that the identity of the female parent accounts for a large proportion of variance in parasite loads. This opens for several possible explanations, including variation in maternal behaviour, but also a potential effect of host genetic factors. Since laying date is highly repeatable and heritable across females in these populations (Caro et al 2009), and considering our results that larval intensity increases with temperature, it is likely that a quantitative genetics analysis of parasite load would reveal heritability of parasite load. Nestbox identity was the second most important random effect after female identity, suggesting that other nest environmental factors might have an effect, and that parasites can be one component of variation in territory quality. Further investigation will be necessary to disentangle the diverse influences of host behavioural, genetic and environmental factors, as well as their implications in terms of selection on both hosts and parasites.

The apparent link between parasite loads and temperature fluctuations in this Mediterranean area is relevant for host-parasite interactions at broader spatial scales. Geographical distributions are expected to be limited by climate, and especially by temperature (David et al. 2003, Kingsolver and Buckley 2017). In a number of insect species the amplitude of thermal tolerance seems reflected in species latitudinal ranges (e.g. Calosi et al. 2010) and in endotherms there is evidence that range limits relate to thermal tolerance limits (e.g. Khaliq et al. 2017). *P. azurea* and *P. falcozi*, the two species present in our study area, are both widespread over the Palaearctic in a wide range of host species (Wesolowski 2001, Matyukhin and Krivosheina 2008). *P. azurea* was reported from Spain to Scandinavia (Potti 2008, Eeva et al. 2015), and *P. falcozi* mostly from Central Europe, but also Germany and Corsica (Wesolowski 2001, Janoskova et al. 2010). Strikingly, of all reports from the Palaearctic it is in Corsica that prevalences and intensities are the highest (Hurtrez-Boussès et al. 1997). This is consistent with the idea that parasite abundances increase with minimal and average spring temperatures, as both are higher on Corsica than in most other locations in Europe. Furthermore *Protocalliphora* abundances relate negatively to average summer temperature; this suggests that they may be limited by heat in areas located further south in the Mediterranean.

Blowfly prevalence and intensities in Western, Central and Northern Europe seem moderate at the moment: prevalence scarcely exceeds 50%, and the reported average intensities remain under or around two larvae per nestling (Wesolowski 2001, Potti 2008, Eeva et al. 2015). However, current projections (IPCC 2013) indicate that regardless of the scenario, by 2100 mean spring and summer temperatures will likely have increased not only in the Mediterranean (spring: +1 °C to +5 °C; summer: +1.5 °C to +7 °C) but also in Central (spring: +1 °C to +5 °C; summer: +2 °C to +6 °C) and Northern Europe (spring: +2 °C to +5 °C; summer: +2 °C to +5 °C). Our results suggest that this might result in modified bird – blowfly interactions. A more comprehensive understanding of the relationships between climate, spring phenology, but also selection and adaptive evolution in bird hosts as well as their parasites is now needed.

## Data accessibility

The data and script used in this preprint are available on Zenodo (https://doi.org/10.5281/zenodo.2576394).

## Ethical statement

Captures were performed under personal ringing permits delivered by the CRBPO (Centre de Recherches par le Baguage des Populations d’Oiseaux) to Anne Charmantier (ringing permit number 1907), Adèle Mennerat, Philippe Perret, and Marcel Lambrechts (permit 1318). All experimental protocols were approved by the ethics committee for animal experimentation of Languedoc Roussillon (305-CEEA-LR- 12066 approved in 2012) as well as by Regional Institutions (bylaw issued by the Prefecture on 15/06/2012 n° 2012167-0003).

## Acknowledgements

We thank the many students and co-workers that helped maintain the study sites for so many years and all the colleagues for discussions or advice. Monitoring of the long-term studies was financially supported by the CNRS, the OSU (OREME), the Mediterranean Centre for Environment and Biodiversity (LabEx CeMEB), and several national or international organizations in the past (European Commission, European METABIRD project, French ANR). The communities that hosted the study plots kindly allowed access to the different (private) study areas. We also wish to thank Fango-MAB and APEEM for logistic support.

This preprint has been reviewed and recommended by *Peer Community In Ecology* (https://dx.doi.org/10.24072/pci.ecology.100012). We are grateful to two anonymous referees for their suggestions that helped improve the quality of this preprint.

## Appendix

**Figure S1.**
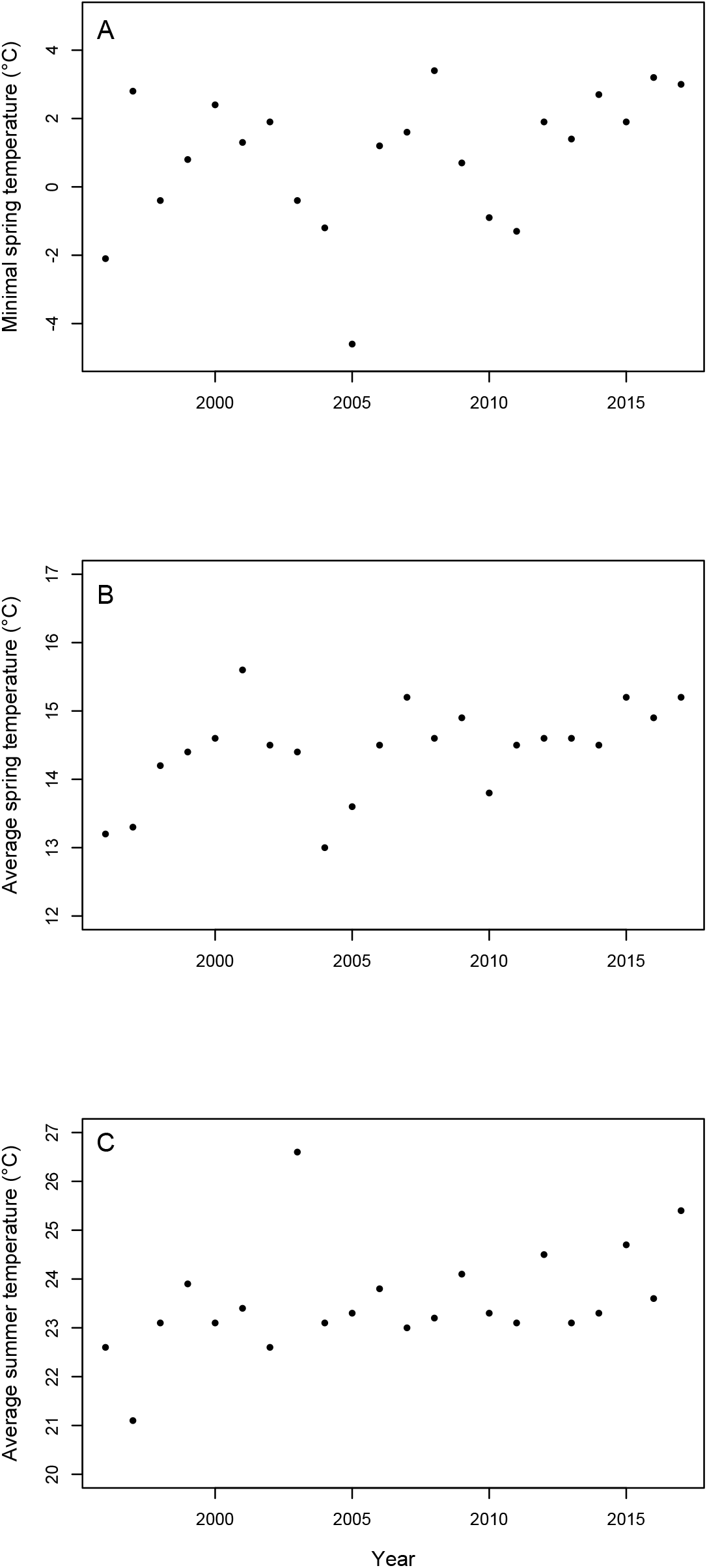
Yearly variation over the 18 years of the study in (A) minimal spring temperature, (B) average spring temperature, and (C) average summer temperature.

**Figure S2.**
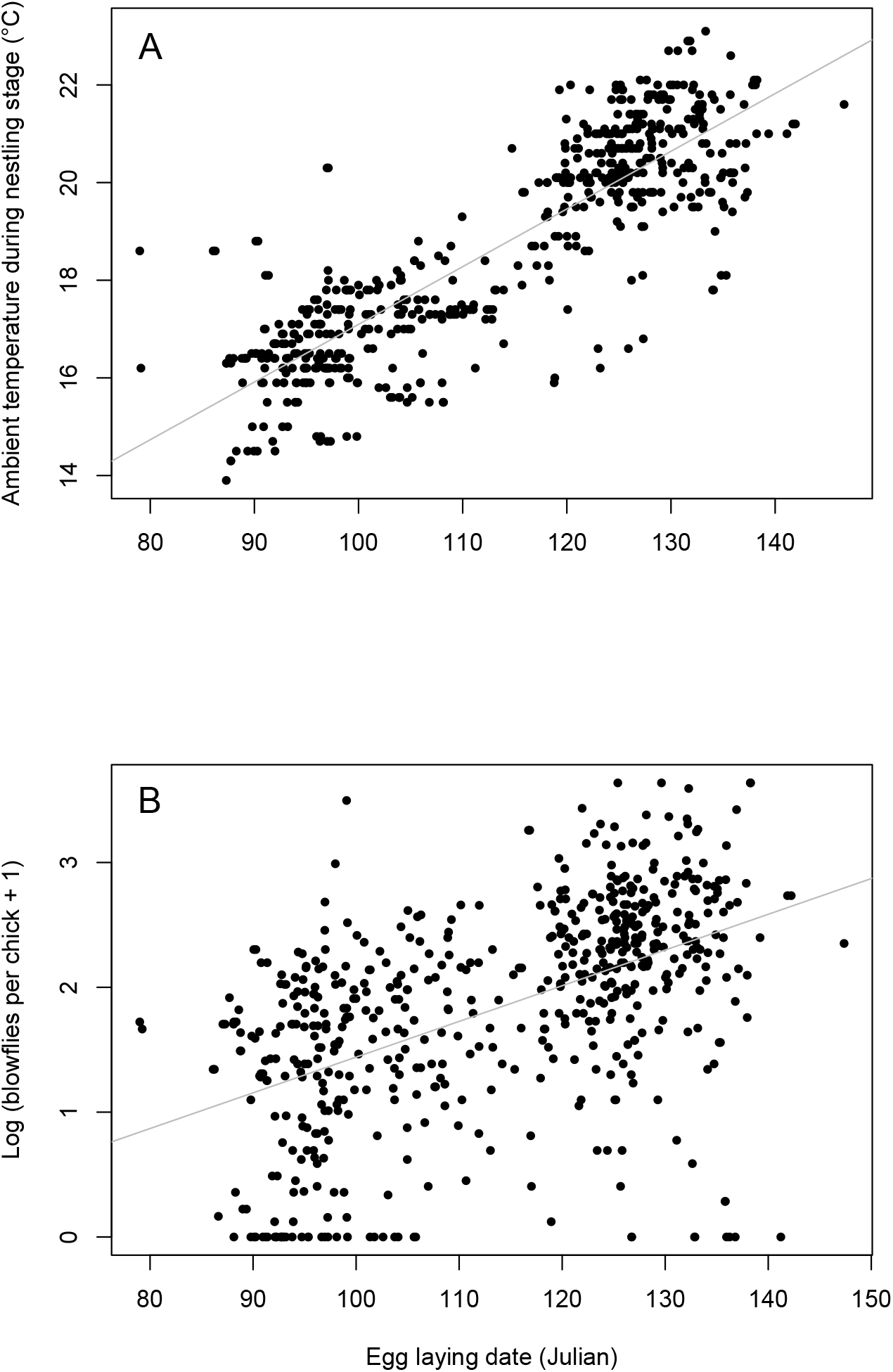
Both average ambient temperature (A) and parasite load (B) during the nestling stage increase with egg-laying date.

## References

Altizer, S. et al. 2013. Climate Change and Infectious Diseases: From Evidence to a Predictive Framework. - Science 341: 514–519.

Andreasson, F. et al. 2018. Experimentally increased nest temperature affects body temperature, growth and apparent survival in blue tit nestlings. - J. Avian Biol. 49: 1–14.

Banbura, J. et al. 2001. Sex differences in parental care in a Corsican Blue Tit *Parus caeruleus* population. - Ardea 89: 517–526.

Bańbura, J. et al. 2004. Effects of *Protocalliphora* Parasites on Nestling Food Composition in Corsican Blue Tits *Parus caeruleus*: Consequences for Nestling Performance. - Acta Ornithol. 39: 93–103.

Bennett, G. and Whitworth, T.. 1991. Studies on the life history of some species of *Protocalliphora* (*Diptera*: *Calliphoridae*). - Can. J. Zool. 69: 2048–2058.

Berrigan, D. et al. 2000. Resistance to temperature extremes between and within life cycle stages in *Drosophila serrata, D. birchii* and their hybrids: intraspecific and interspecific comparisons. - Biol. J. Linn. Soc.: 403–416.

Blondel, J. 1985. Breeding Strategies of the Blue Tit and Coal Tit (*Parus*) in Mainland and Island Mediterranean Habitats: A Comparison. - J. Anim. Ecol. 54: 531–556.

Blondel, J. et al. 2006. A Thirty-Year Study of Phenotypic and Genetic Variation of Blue Tits in Mediterranean Habitat Mosaics. - Bioscience 56: 661.

Brooks, D. R. and Hoberg, E. P. 2007. How will global climate change affect parasite-host assemblages? - Trends Parasitol. 23: 571–574.

Calosi, P. et al. 2010. What determines a species’ geographical range? Thermal biology and latitudinal range size relationships in European diving beetles (*Coleoptera: Dytiscidae*). - J. Anim. Ecol. 79: 194–204.

Cantarero, A. et al. 2013. Behavioural responses to ectoparasites in pied flycatchers *Ficedula hypoleuca*: An experimental study. - J. Avian Biol. 44: 591–599.

Castaño-Vázquez, F. et al. 2018. Experimental manipulation of temperature reduce ectoparasites in nests of blue tits *Cyanistes caeruleus*. - J. Avian Biol. 49: 1–6.

Charlier, J. et al. 2016. Climate-driven longitudinal trends in pasture-borne helminth infections of dairy cattle. - Int. J. Parasitol. 46: 1–8.

Charmantier, A. et al. 2004. Parasitism reduces the potential for evolution in a wild bird population. - Evolution 58: 203–206.

Chown, S. and Nicolson, S. 2004. Insect physiological ecology. - Oxford University Press.

Christe, P. et al. 1994. Ectoparasite affects choice and use of roost sites in the great tit, *Parus major*. - Anim. Behav.: 895–898.

1997. Host parasite evolution: general principles and avian models (D Clayton and J Moore, Eds.). - Oxford University Press.

David, J. R. et al. 2003. The fly that came in from the cold : geographic variation time from low-temperature of recovery exposure in Drosophila subobscura. 17: 425–430.

Dawson, R. D. et al. 2005. Effects of Experimental Variation in Temperature on Larval Densities of Parasitic *Protocalliphora* (*Diptera* : *Calliphoridae*) in Nests of Tree Swallows (*Passeriformes* : *Hirundinidae*). - Physiol. Ecol. 34: 563–568.

Descamps, S. et al. 2002. Asynchronous hatching in a blue tit population: a test of some predictions related to ectoparasites. - Can. J. Zool. 80: 1480–1484.

Dudaniec, R. Y. et al. 2007. Interannual and interspecific variation in intensity of the parasitic fly, *Philornis downsi*, in Darwin’s finches. - Biol. Conserv. 139: 325–332.

Eads, D. and Hoogland, J. 2016. Factors that affect parasitism of black-tailed prairie dogs by fleas. - Ecosphere 7: e01372.

Eeva, T. L. E. and Nurmi, J. 1994. Effects of ectoparasites on breeding success of great tits (*Parus major*) and pied flycatchers (*Ficedula hypoleuca*) in an air pollution gradient. - Can. J. Zool. 72: 624–635.

Eeva, T. et al. 2015. Species and abundance of ectoparasitic flies (*Diptera*) in pied flycatcher nests in Fennoscandia. - Parasites and Vectors 8: 1–9.

Elderd, B. D. and Reilly, J. R. 2014. Warmer temperatures increase disease transmission and outbreak intensity in a host-pathogen system. - J. Anim. Ecol. 83: 838–849.

Enriquez, T. and Colinet, H. 2017. Basal tolerance to heat and cold exposure of the spotted wing drosophila, *Drosophila suzukii*. - PeerJ 5: e3112.

Feder, M. E. et al. 1997. Natural thermal stress and heat-shock protein expression in *Drosophila* larvae and pupae. - Funct Ecol 11: 90–100.

Grosbois, V. et al. 2006. Climate impacts on Mediterranean blue tit survival: an investigation across seasons and spatial scales. - Glob. Chang. Biol. 12: 2235–2249.

Harvell, C. et al. 2002. Climate Warming and Disease Risks for Terrestrial and Marine Biota. - Science 296: 2158–2163.

Heeb, P. et al. 1996. Horizontal Transmission and Reproductive Rates of Hen Fleas in Great Tit Nests. - J. Anim. Ecol. 65: 474–484.

Heeb, P. et al. 2000. Bird–Ectoparasite Interactions, Nest Humidity, and Ectoparasite Community Structure. - Ecology 81: 958–968.

Hernandez, A. D. et al. 2013. Climate changes influence free-living stages of soil-transmitted parasites of European rabbits. - Glob. Chang. Biol. 19: 1028–42.

Hudson, P. J. et al. 2006. Is a healthy ecosystem one that is rich in parasites? - Trends Ecol. Evol. 21: 381–5.

Hurtrez-Boussès, S. et al. 1997. High blowfly parasitic loads affect breeding success in a Mediterranean population of blue tits. - Oecologia: 514–517.

Hurtrez-Boussès, S. et al. 1998. Chick parasitism by blowflies affects feeding rates in a Mediterranean population of blue tits. - Ecol. Lett. 1: 17–20.

Hurtrez-Boussès, S. et al. 1999. Variations in prevalence and intensity of blow fly infestations in an insular Mediterranean population of blue tits. - Can. J. Zool. 77: 337–341.

Hurtrez-Boussès, S. et al. 2000. Effects of ectoparasites of young on parents’ behaviour in a Mediterranean population of Blue Tits. - J. Avian Biol. 31: 266–269.

IPCC 2013. Annex I: Atlas of Global and Regional Climate Projections. - In: van Oldenborgh, G. J. et al. (eds), Climate Change 2013: The Physical Science Basis. Contribution of Working Group I to the Fifth Assessment Report of the Intergovernmental Panel on Climate Change. Cambridge University Press, in press.

Jamriska, J. et al. 2010. Host spectrum of bird blow flies of the genus Protocalliphora Hough, 1899 (Diptera, Calliphoridae) in Slovakia. - Sylvia 46: 125–132.

Janoskova, V. et al. 2010. Pre-imaginal stages of the blowfly *Protocalliphora falcozi* in nests of the tree sparrow (Passer montanus). - Entomol. Fenn. 21: 107–116.

Khaliq, I. et al. 2017. The influence of thermal tolerances on geographical ranges of endotherms. - Glob. Ecol. Biogeogr. 26: 650–668.

Kingsolver, J. G. and Buckley, L. B. 2017. Quantifying thermal extremes and biological variation to predict evolutionary responses to changing climate. - Philos. Trans. R. Soc. B Biol. Sci. 372: 20160147.

Kleindorfer, S. and Dudaniec, R. Y. 2009. Love thy neighbour? Social nesting pattern, host mass and nest size affect ectoparasite intensity in Darwin’s tree finches. - Behav. Ecol. Sociobiol. 63: 731–739.

Lambrechts, M. M. et al. 2004. Habitat quality as a predictor of spatial variation in blue tit reproductive performance: A multi-plot analysis in a heterogeneous landscape. - Oecologia 141: 555–561.

Loye, J. and Zuk, M. 1991. Bird - parasite interactions: ecology, evolution and behaviour. - Oxford University Press.

Marrot, P. et al. 2017. Multiple extreme climatic events strengthen selection for earlier breeding in a wild passerine. - Philos Trans R Soc L. B Biol Sci 372: 20160372.

Marrot, P. et al. 2018. Current spring warming as a driver of selection on reproductive timing in a wild passerine. - J. Anim. Ecol. 87: 754–764.

Mas-Coma, S. et al. 2009. Climate change effects on trematodiases, with emphasis on zoonotic fascioliasis and schistosomiasis. - Vet. Parasitol. 163: 264–280.

Matyukhin, A. and Krivosheina, M. 2008. Contribution to the knowledge of *Diptera* (*Insecta*) parasitizing on birds. - Entomol. Rev. 88: 258–259.

Mennerat, A. et al. 2008. Aromatic plants in blue tit *Cyanistes caeruleus* nests: no negative effect on blood-sucking *Protocalliphora* blow fly larvae. - J. Avian Biol. 39: 127–132.

Mennerat, A. et al. 2009. Aromatic plants in nests of the blue tit *Cyanistes caeruleus* protect chicks from bacteria. - Oecologia 161: 849–855.

Merino, S. and Potti, J. 1996. Weather dependent effects of nest ectoparasites on their bird hosts. - Ecography (Cop.). 19: 107–113.

Møller, A. et al. 1990. Fitness effects of parasites on passerine birds: a review. - In: Blondel, J. et al. (ed), Population biology of passerine birds. Springer-Verlag, pp. 269–280.

Møller, A. P. et al. 2013. Assessing the effects of climate on host-parasite interactions: A comparative study of european birds and their parasites. - PLoS One in press.

Moreno, J. et al. 2009. Nest-dwelling ectoparasites of two sympatric hole-nesting passerines in relation to nest composition: An experimental study. - Écoscience 16: 418–427.

Ogden, N. H. and Lindsay, L. R. 2016. Effects of Climate and Climate Change on Vectors and Vector-Borne Diseases: Ticks Are Different. - Trends Parasitol. 32: 646–656.

Owen, D. and Ash, J. 1955. Additional records of *Protocalliphora* (*Diptera*) in birds’ nests. - Br. Birds XLVIII: 225–229.

Porlier, M. et al. 2012. Variation in phenotypic plasticity and selection patterns in blue tit breeding time: Between- and within-population comparisons. - J. Anim. Ecol. 81: 1041–1051.

Potti, J. 2008. Blowfly Infestation at the Nestling Stage Affects Egg Size in the Pied Flycatcher *Ficedula hypoleuca*. - Acta Ornithol. 43: 76–82.

Remeš, V. and Krist, M. 2005. Nest design and the abundance of parasitic *Protocalliphora* blow flies in two hole-nesting passerines. - Ecoscience 12: 549–553.

Rohr, J. R. et al. 2011. Frontiers in climate change-disease research. - Trends Ecol. Evol. 26: 270–277.

Roiz, D. et al. 2014. Climatic effects on mosquito abundance in Mediterranean wetlands. - Parasit. Vectors 7: 333.

Rose, H. et al. 2014. Exploiting parallels between livestock and wildlife: Predicting the impact of climate change on gastrointestinal nematodes in ruminants. - Int. J. Parasitol. Parasites Wildl. 3: 209–219.

Simon, A. et al. 2004. Physiological ecology of Mediterranean blue tits (*Parus caeruleus* L.): effects of ectoparasites (*Protocalliphora* spp.) and food abundance on metabolic capacity of nestlings. - Physiol. Biochem. Zool. 77: 492–501.

Simon, A. et al. 2005. Impact of Ectoparasitic Blowfly Larvae (*Protocalliphora* Spp.) on the Behavior and Energetics of Nestling Blue Tits. - J. F. Ornithol. 76: 402–410.

Stromberg, B. E. 1997. Environmental factors influencing transmission. - Vet. Parasitol. 72: 247–264.

Thomas, D. et al. 2007. Common paths link food abundance and ectoparasite loads to physiological performance and recruitment in nestling blue tits. - Funct. Ecol. 21: 947–955.

van Noordwijk, A. J. et al. 1995. Selection for the Timing of Great Tit Breeding in Relation to Caterpillar Growth and Temperature. - J. Anim. Ecol. 64: 451.

Visser, M. E. et al. 2006. Shifts in Caterpillar Biomass Phenology Due to Climate Change and Its Impact on the Breeding Biology of an Insectivorous Bird. - Oecologia 147: 164–172.

Wesolowski, T. 2001. Host - parasite interactions in natural holes: marsh tits (*Parus palustris*) and blow flies (*Protocalliphora falcozi*). - J. Zool.: 495–503.

